# A minimal biophysical model recapitulates the maintenance of whole-genome histone methylation patterns across cell division

**DOI:** 10.64898/2026.01.16.699330

**Authors:** Erik J. Navarro, William Poole, Ariane Lismer, Jieqiong Qu, Ioannis Kafetzopoulos, Aled Parry, Jannat Ijaz, Alice Santambrogio, Ryan Spangler, Francisco M. Martín-Zamora, Ken Raj, Wolf Reik, Hana El-Samad, Carlos F. Lopez, Simone Bianco

**Affiliations:** Altos Labs - Institute of Computation; Altos Labs - Institute of Science; Altos Labs - Institute of Technology

**Keywords:** biomolecular condensates, epigenetic inheritance, chromatin biophysics

## Abstract

In multi-cellular eukaryotic organisms, cell type and specific functional identity are defined by the epigenetic patterning of chemical modifications to DNA and chromatin that modulate the expression and silencing of specific genes. When a cell divides, histones containing important epigenetic marks are distributed between the two daughter strands leading to a temporary dilution of epigenetic information and cell identity. In this work we introduce a physics-based model of epigenetic memory that explains how cells restore and maintain H3K9me3 and H3K27me3 histone methylation patterning after cell division. We demonstrate that emergence and maintenance of the epigenetic program can be explained by a minimal mechanism that makes use of the biophysics of polymers, phase condensates and enzymatic activity. Because polymer looping and enzyme co-localization are generic, ubiquitous properties of chromatin, our model shows that a baseline of epigenetic memory can be maintained and inherited genome-wide without invoking sequence-or location-specific regulatory mechanisms. We validate our model via genome-wide epigenetic time-course simulation and comparison to experimental epigenetic data from multiple donors, multiple cell types, and for multiple epigenetic marks. Finally, we use our model as a conceptual framework to understand cellular reprogramming by hypothesizing that these processes first contend with and later utilize epigenetic maintenance programs.

**Significance Statement:** Epigenetic patterns—chemical modifications of DNA and histones—are central to establishing cell identity and are implicated in many diseases. However, the mechanisms of epigenetic pattern maintenance are not fully understood. We introduce the Looping, Enzyme, and Phase-Condensate (LEAP) model of H3K9me3 and H3K27me3 epigenetic pattern restoration. LEAP shows that generic chromatin biophysics—the statistical physics of polymer looping and mark-dependent methyltransferase co-localization—suffices for genome-wide epigenetic memory, without sequence-or location-specific elements. We quantitatively validate LEAP by simulating the whole genome and comparing it against diverse human epigenetic data. LEAP maintains patterns at the scale of topologically associating domains and sets a baseline on which more complex mechanisms act, providing a framework for epigenetic aging and reprogramming.

## 1 Introduction

Chromatin, the highly dynamic molecular complex composed of DNA, RNA, and proteins, is involved in many aspects of cellular behavior, ranging from adaptation and response to perturbations to organismal physiology (*1*). Epigenetics modulates the expression of genes through physically and chemically modifying chromatin in a cell-type specific manner patterned over the course of development (*2*). The progressive loss of epigenetic stability has been linked to aging and disease, and recent evidence has shown that the epigenome is reset during partial cellular reprogramming and rejuvenation (*3*).

Among the epigenetic modifications that occur during development, histone modifications are of particular importance. Histones are the proteins that bind to DNA forming molecular octamers called nucleosomes, which are one of the main structural elements of chromatin. Histones are frequently chemically modified via a complex code of acetylation, methylation, ubiquitination, and phosphorylation marks which help determine cell type by modulating gene expression (*4*). Two chromatin-silencing histone methylation marks are known to be of particular importance in determining cell state: H3K9me3 and H3K27me3, which are catalyzed (in humans) by methyltransferases SUV39 and PRC2, respectively (*5*). Their loss is directly linked to aging and results in chromatin instability and increased cellular stress (*6*). Additionally, partial reprogramming has been associated with restoration of epigenetic marks (*7*).

Unlike the inheritance of genetic information encoded as a DNA sequence, histone marks like H3K9me3 and H3K27me3 are not directly copied during DNA replication. In humans it has been established that modified parental histones bearing H3K9me3 and H3K27me3 modifications are distributed symmetrically (meaning randomly with equal probability) between the two daughter cells during cell division (*8*). This means that after division, each daughter cell receives a random combination of epigenetically modified histones and newly synthesized naive histones not yet bearing epigenetic modifications (*8, 9*). The daughter cells are then tasked with restoring the full epigenetic pattern from this diluted signal in order to preserve cellular identity and function. On the other hand, full and partial reprogramming aim to rewrite and restore the epigenome, respectively. It is unlikely that partial or full reprogramming via Yamanaka factors fundamentally changes the mechanisms of epigenetic inheritance. Therefore, the formation and maintenance of epigenetic memory during cellular division is not merely a characteristic of cellular health, but a fundamental process that is both utilized and challenged by cellular reprogramming.

The mechanisms of epigenetic inheritance, stability, and memory across generations remain major outstanding questions in cellular biology, and a rich body of theory has proposed how such patterns might be established and maintained. Rather than a single mechanism, this body of work has introduced a range of molecular features that individual models combine in different subsets, which we group into three broad classes. The first invokes sequence-specific genomic elements: a dedicated nucleation site from which marks spread (*10–12*), boundary or insulator elements that delimit domains (*13, 14*), and locus-specific kinetic parameters (*15*). The second relies on interplay between distinct epigenetic species: bistable switching between mutually antagonistic marks (*13, 14*), or cross-talk between histone marks and DNA methylation (*16*). The third emphasizes 3D polymer dynamics coupled with biochemical interactions: looping brings distant loci into contact (*11, 12*), co-localization of methyltransferases with methylated histones and a limiting enzyme pool concentrate catalytic activity (*15, 17, 18*), and attractive forces between methylated histones, together with chromatin compaction during mitosis, stabilize the resulting domains (*18*). These features are not mutually exclusive—most models combine several, such as nucleation with looping (*11*) or enzyme limitation with compaction (*15, 18*)—and together they form the foundation on which our work builds. Each feature, however, is a specialized ingredient that need not be present at every epigenetic domain, and these mechanisms have largely been examined in idealized or small-scale settings, with little genome-wide validation using modalities such as ChIP-seq or CUT&RUN; one notable exception achieves fewer than 10 generations of pattern maintenance (*17*) on such a validation. Table S1 summarizes the features invoked by these models and the scope of their validation.

Our work builds on these efforts to model the inheritance and restoration of H3K9me3 and H3K27me3 methylation marks through the cell cycle as illustrated in Figure 1 A. We propose a much simplified model, which we call the Looping, Enzyme And Phase-Condensate (LEAP) model, involving polymer looping mediated catalysis and co-localization of methyltransferases to densely methylated genomic regions. We show that this minimal mechanism is sufficient to generate and maintain epigenetic patterns that persist across many cell generations. Because it relies only on polymer looping and enzyme co-localization—generic, ubiquitous properties of chromatin—LEAP shows that a baseline of epigenetic memory can be established and inherited without sequence-or location-specific regulatory mechanisms, or large differences in chromatin density between domains, whose existence has recently been questioned (*19, 20*). We conceptualize LEAP as a foundation upon which other biological mechanisms, such as aging and cellular reprogramming, can act to further regulate epigenetic patterns. We validate this model on genome-scale H3K9me3 and H3K27me3 data from multiple donors and cell types.

**Figure 1:**
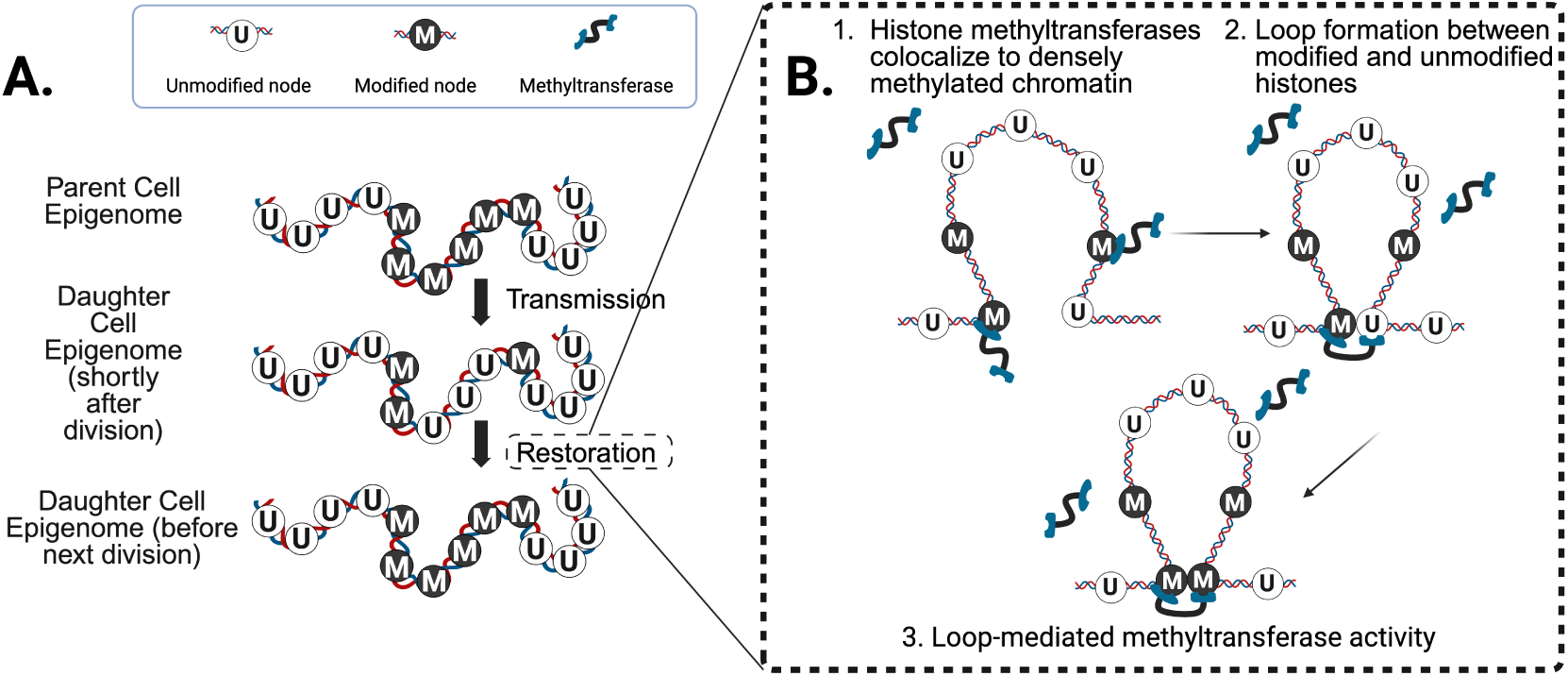
A. Cartoon illustration of epigenetic dilution due to cell-division and subsequent restoration. B. A cartoon of the proposed mechanism of epigenetic restoration; (1) histone methyltransferase co-localization to methylated chromatin, (2) loop formation, and (3) loop mediated catalysis.

## 2 The Looping, Enzyme And Phase-condensate (LEAP) Model

### Conceptual Model

The Looping, enzyme, and phase-condensate (LEAP) model of epigenetic restoration combines three experimental observations about histone methyltransferases. First, these enzymes are known to bind at higher frequency to their own reaction product. This implies positive feedback or a strong preference for the enzymes to interact with the methylated state rather than the unmethylated or other state (*21, 22*). This reader-writer capacity was first established for the H3K9 methyltransferase Clr4/Suv39 (*23*). Second, these enzymes’ catalytic activity is mediated by cooperatively binding to a methylated histone and an unmethylated histone simultaneously (*21, 22*). Essentially, by binding to its own product, the histone methyltransferase can methylate adjacent histones and ensure that the mark is maintained, reinforced or potentially spread across the local chromatin structure. It is well known that polymers in solution, including chromatin, produce loops which bring together linearly separated regions of DNA. The observed probability of two monomers coming into contact scales like an inverse power law with the linear distance and an exponent that depends on the solvent conditions and may vary between cell and organism types (*24*).

Third, it has been observed that H3K9me3 and H3K27me3 methyltransferases co-localize in high density condensates that form around regions of dense H3K9 and H3K27 methylation (*25, 26*). Consistent with a density-dependent mechanism, efficient spreading and inheritance of H3K9 methylation requires a critical density of the mark (*27*). The formation of these condensates potentially increases the local reaction rate and ensures epigenetic specificity by enforcing a spatial organization. After cell division, even though only half the original number of methylated histones are present, there is still sufficient density for enzyme condensates to localize around epigenetic domains. These regions are then able to reliably restore themselves due to polymer-polymer looping which ensures that methylation is contained in a relatively local region around these dense condensates. This model is illustrated in Figure 1 B.

### Idealized Model

We mathematically model an idealized epigenetic code consisting of a single epigenetic mark (methylation) which resides along the monomers of a polymer chain representing chromatin. This polymer has *N* monomers (nucleosomes containing a modifiable histone), each denoted by *x_i_* ∈ {0, 1}, which indicates whether the histone *i* is unmodified (*x_i_* = 0) or methylated (*x_i_* = 1). The ordering of these monomers in a linear chain *X* = (*x*_1_, …, *x_i_*, …, *x_N_*) constitutes a specific epigenetic pattern. The pattern degrades due to dilution at cell division, modeled as binomial partitioning occurring regularly with a frequency 1/*t_div_*. The pattern restores itself via loop-mediated methyltransferase activity. Loops are allowed to form between sites *i* and *j* with probability *P_L_* through the polymer physics derived inverse-power law (*28*). Methyltransferases (*E*) may be bound to any given methylated monomer with probability *P_E_* modeled as a sigmoid function of the local marked-monomer density along the chain, representing the non-linear effects of methyltransferase phase condensation and co-localization to methylated histones. The resultant rate at which a monomer *i* may gain a mark, *Q*(*x_i_* = 0 → *x_i_* = 1), is given by:

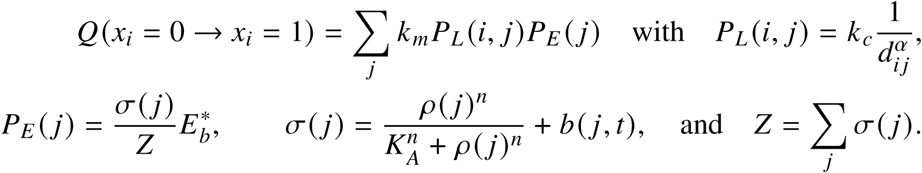

Here *d_i j_* is the distance between monomers *i* and *j* along the polymer chain and *b*(*j*, *t*) is a time-dependent bias term assumed to be 0 unless otherwise noted. The local monomer density *ρ*(*j*) is computed with a Gaussian kernel density estimate with bandwidth *λ*. Additionally, *t_div_* is the division time, *k_m_* is the methylation rate of the methyltransferase, *k_c_* is the contact rate related to polymer mobility, *α* is the scaling exponent of the contact law, *K_A_* and *n* parameterize the non-linearity *σ*, and 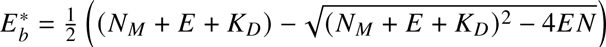 is the total number of bound enzymes which depends on the total number of enzymes *E*, the dissociation constant *K_D_*, and the number of marked monomers *N_M_* = Σ*_i_x_i_*. This model is formalized and simulated as a continuous time Markov chain (CTMC) as elaborated in the SI A.

### Coarse Grained Model

Simulating the above model at the genomic scale is prohibitively expensive from a computational standpoint due to the quadratic number of terms required to evaluate *Q*(*x_i_* = 0 → *x_i_* = 1) for all *i*. The human genome has around 30 × 10^6^ nucleosomes and larger chromosomes may have over 10^6^ nucleosomes which would result in an equation with ∼ 10^12^ terms evaluated at every simulation timestep. In order to carry out genome-scale simulations, we developed a coarse grained model which allows us to simulate sets of marks pooled together into a single monomer. Briefly, instead of simulating dynamics for *X* ∈ {0, 1}*^N^* where each monomer represents a single marked or unmarked position of chromatin, we simulate *Y* ∈ {0, 1, …*μ*}*^ν^* where *μ* is the maximum number of marks at any given position and *ν* = *N*/*μ* is the number of coarse grained monomers. For example, *μ* = 2 would be appropriate for a nucleosome-level simulation where each nucleosome can have 0, 1, or 2 marks. If *μ* > 2, this indicates that multiple nucleosomes are coarse-grained into a single simulated monomer. For a detailed description of this model and simulation details see SI C.

## 3 Results

### 3.1 The mechanisms captured by the idealized LEAP model are sufficient to generate epigenetic memory

We show that epigenetic patterns can be stably maintained over hundreds of cell cycles using an illustrative pattern simulated with the idealized LEAP model. These dynamics are visualized over a single generation, multiple generations, and as transition rates in Figure 2 A, B, and D. We quantify epigenetic drift by computing the auto-correlation between the initial pattern and later patterns in Figure 2 C. Notice that this correlation is much higher when the data is smoothed because the stochastic nature of the model results in high variance at the single mark scale despite epigenetic domains being largely stable.

**Figure 2:**
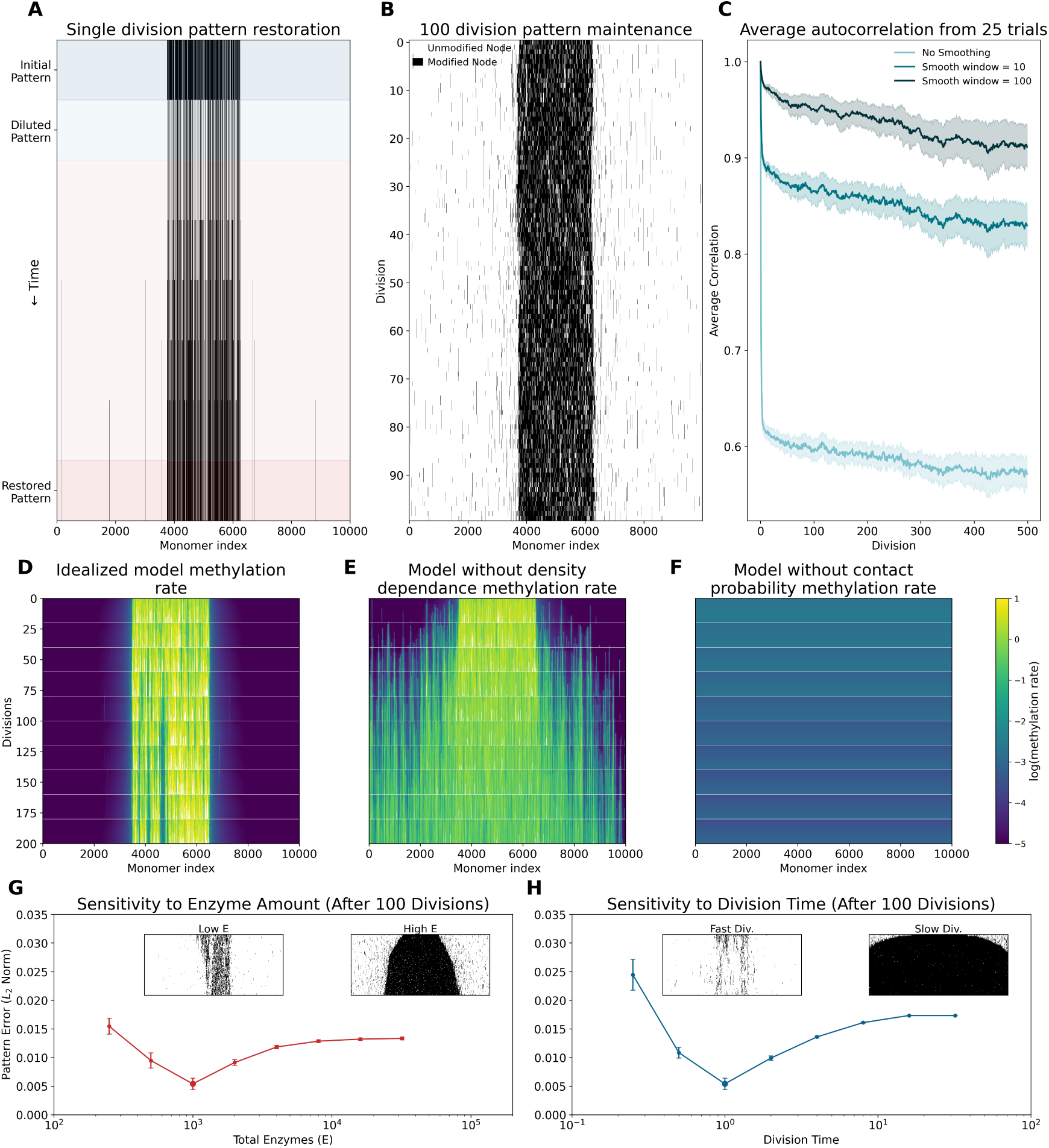
A. Simulation results for a single dilution-restoration cycle. B. Simulation results for 100 dilution-restoration cycles. C. Auto-correlation computed between the initial condition and subsequent time-points; correlations are computed using data smoothed with a uniform window. D. The methylation rate simulated with the idealized model. E. The methylation rate simulated using a model without a density dependence term. F. the methylation rate simulated using a model without the contact probability term. G. Idealized model sensitivity to the enzyme amount. H. Idealized model sensitivity to the division time.

The simulations demonstrate that polymer looping and methyltransferase methyl-mark co-condensate formation are both necessary to stably maintain epigenetic patterns. The co-condensate formation provides a non-linearity that stabilizes the mark locally and dramatically diminishes spreading but is alone insufficient to localize patterns. Removing this methylation density dependence non-linearity *σ* weakens the spatial localization of the pattern and allows methylation to diffuse outwards due to the 1/*d^α^* contact probability (Figure 2E). The contact probability term allows for pattern localization along the chain but cannot constrain them from rapidly spreading. Removing the inverse power law contact dependence *P_L_* results in a uniform methylation rate across the chain with no spatial structure (Figure 2F). Furthermore, these two features are together sufficient for spontaneous pattern formation: starting from random initial conditions, the model reliably forms well-defined, metastable domains, as investigated in SI B. Finally, the sensitivity of the maintained patterns to the total enzyme amount *E* and to the division time is shown in Figure 2 G and H, and is discussed below.

### 3.2 The coarse-grained LEAP model shows remarkable accuracy in predicting epigenetic marks for hundreds of generations in biological experiments, across different ages and cell types

We apply the coarse LEAP model with *μ* = 10^4^, corresponding to ∼50 nucleosomes per coarse grained monomer, to simulate real H3K9me3 and H3K27me3 data collected from passaged HU-VECs and human derived fibroblasts (HDFs), see SI G and G.4. Example snapshots and simulated dynamics of the model can be seen in the first two columns of Figure 3 (panels A, B, E, F, I, and J). This model is applied to HDF H3K27me3 (first row, Figure 3A–D), HDF H3K9me3 (second row, Figure 3E–H), and HUVEC H3K27me3 (third row, Figure 3I–L) data. Additional simulation output and implementation details can be found in SI D. Notably, the model is able to recapitulate H3K9me3 and H3K27me3 epigenetic patterns in 8 different HDFs for over 100 generations with high accuracy, as measured via the correlation between the initial pattern and the simulated data after 100 generations (Figure 3 D and H). We further compute the auto-correlation times for each mark in each cell type (Figure 3 C, G, K) using both a smoothed (uniform smoothing with a window of 100 coarse monomers which is equivalent to 1Mb) and unsmoothed state to compute the correlation. The increased auto-correlation time of the smoothed data indicates that larger epigenetic patterns on the scale of Mb (which correspond to topologically associating domains (TADs) (*29*)) can be accurately maintained over hundreds of generations. TADs are thought to constrain and control interactions between enhancers and their target genes, and their disruption is linked with loss of cell type specific epigenetic memory (*30*). Finally, due to having time-course epigenetic data from HUVEC cells as a function of cumulative population doubling, we compare the model dynamics to the epigenetic dynamics. We find that our model predicts more epigenetic drift in HUVECs than we see in the actual data illustrated in Figure 3 L by the observation that young simulations (which have not drifted far from data derived from young cells) correlate more strongly to data derived from old passaged cells than old simulations. We conceptualize this discrepancy by noting that our model provides a baseline for epigenetic drift but does not take into account any other biological processes which may be actively maintaining and modulating histone methylation patterns as cells age. Indeed, HUVECs are a common in vitro model of aging in part because they have a well-characterized senescence phenotype that emerges over cell passages (*31*); this cell state transition may be responsible for discrepancies with the LEAP model.

**Figure 3:**
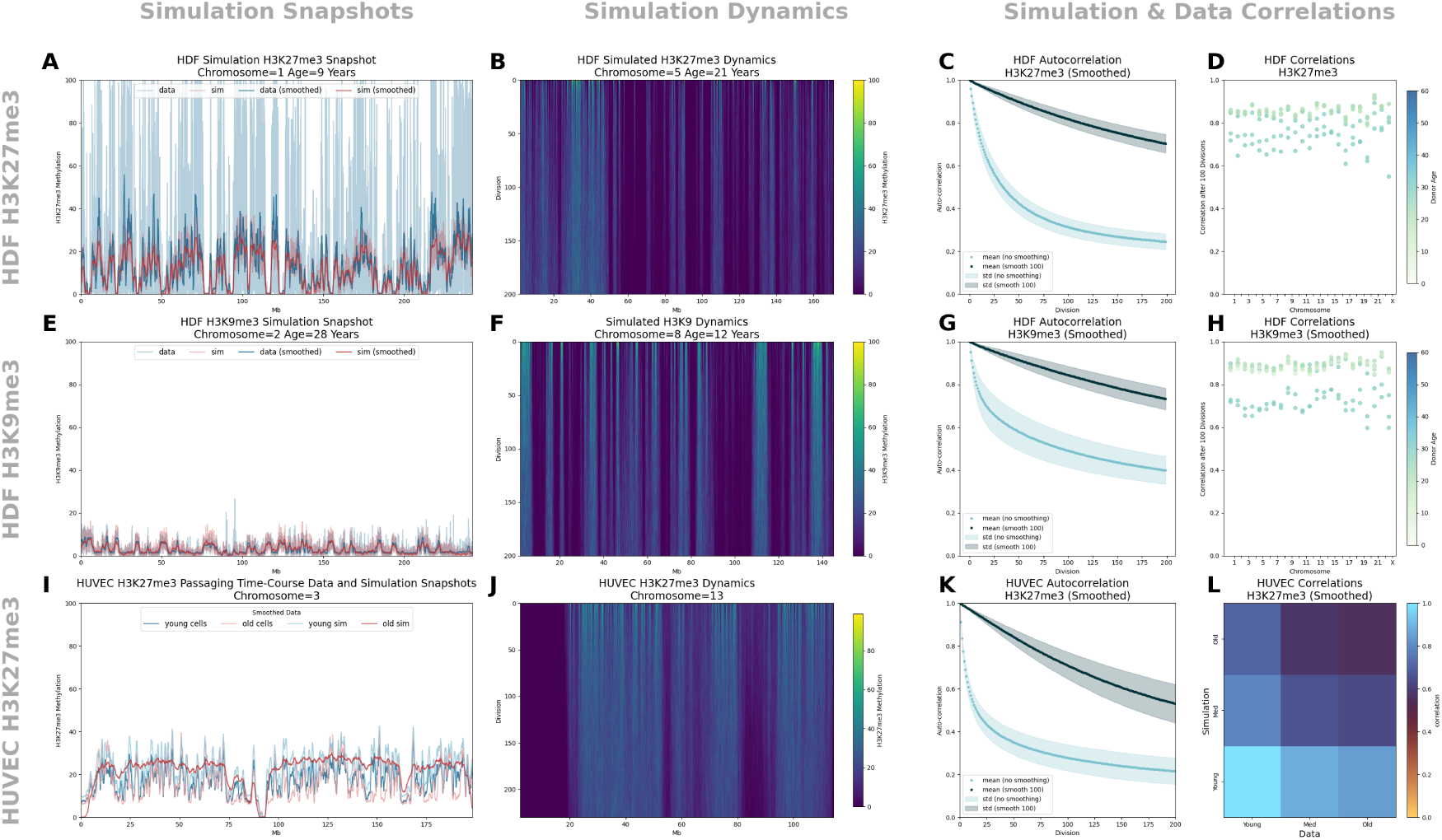
A. Example H3K27me3 data and simulation output after 100 divisions for a single HDF donor and chromosome. B. Example H3K27me3 simulation dynamics for a single HDF donor and chromosome. C. Autocorrelation of the HDF H3K27me3 simulations averaged across all donors and chromosomes shown for unsmoothed data and smoothed data (uniform smoothing with a window of 100). D. Correlation between HDF H3K27me3 smoothed simulation output after 100 generations and smoothed data split by donor and chromosome and colored by age. E. Example H3K9me3 data and simulation output after 100 divisions for a single HDF donor and chromosome. F. Example H3K9me3 simulation dynamics for a single HDF donor and chromosome. G. Auto-correlation of the HDF H3K9me3 simulations averaged across all donors and chromosomes shown for unsmoothed data and smoothed data (uniform smoothing with a window of 100). H. Correlation between HDF H3K9me3 smoothed simulation output after 100 generations and smoothed data split by donor and chromosome and colored by age. I. Young (CPD = 34, simulated divisions = 10), medium (CPD = 150, simulated divisions = 150), and old (CPD = 263, simulated divisions = 225) H3K27me3 data and simulation output. CPD is the cumulative population doubling score. J. Example H3K27me3 simulation dynamics for a single HUVEC chromosome beginning at a young state. K. HUVEC H3K27me3 autocorrelation of the simulation dynamics with the initial state for 25 replicates. L. Correlation between young, medium, and old simulations and data (as described in the caption for panel I).

### 3.3 Insights into the therapeutic effects of reprogramming

To represent the remodeling of the epigenome during development and reprogramming processes (*32*), we use the idealized model with a bias term *b*(*j*, *t*) ≥ 0. The bias term can phenomenologically model therapeutics by varying the dosage (bias value) and application regimen (bias intervals). Both development and cellular reprogramming are known to involve epigenetic changes in H3K9me3 and H3K27me3 marks (*33, 34*). The bias term can be interpreted as a change in the density of methyltransferases at specific sites. This effect mimics important observations of transcription factor mediated reprogramming. It has been observed that reprogramming factors (particularly Oct4) cause reorganization of phase condensates via H3K27 acetylation (*35*), which is mutually exclusive to H3K27me3 (*36*). Moreover, pioneering transcription factors like OSKM are known to recruit chromatin modifying enzymes to the site of binding, effectively biasing the density of enzymes across the genome. In general, our model predicts that any epigenetic modification due to stress, disease, or intervention, may persist in the absence of post-modification correction. Therefore, we use simulations of the idealized model to conceptualize how epigenetic stress (red shaded regions in Figure 4) can create changes in epigenetic patterns that are propagated after the stress is removed, which has been observed experimentally (*33*). Similarly, therapeutic strategies such as rejuvenation via partial reprogramming (*7*) can be seen as epigenetic modifications that restore patterns to a healthier state, which has also been observed via tools such as epigenetic clocks (*37*). Figure 4A shows a healthy baseline in the absence of stress or reprogramming, while Figure 4B–D show how different dosages of epigenetic therapies can result in different outcomes ranging from failure to rejuvenate (continuance of the stress-induced epigenetic pattern) to restoration of the initial epigenetic pattern (e.g. partial reprogramming) to full reprogramming that changes cell identity and results in a qualitatively different epigenetic pattern. Figure 4E shows the bias terms *b*(*j*, *t*) applied in these simulations, and Figure 4F quantifies the correlation of each intervention with the healthy baseline as a function of cell divisions.

**Figure 4:**
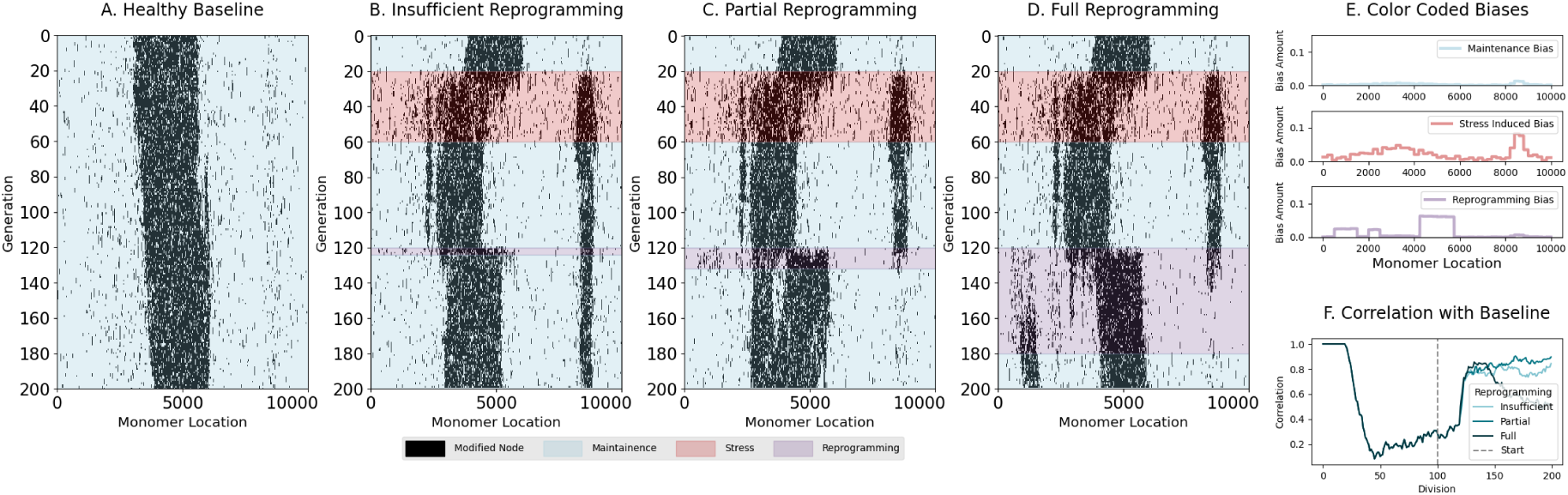
A. Idealized model simulation of a healthy baseline without stress or reprogramming. B. Idealized model simulation with stress and a short reprogramming intervention. C. Idealized model simulation with stress and medium length reprogramming intervention. D. Idealized model simulation with stress and a long reprogramming intervention. E. Bias terms used for the maintenance, stress, and reprogramming periods of the simulation. F. Correlation of the different reprogramming interventions with the healthy baseline as a function of cell divisions using smoothed output (window = 100).

## 4 Discussion

The LEAP model illustrates how the fundamental biophysics of polymer looping, enzyme kinetics, and phase condensate formation can underlie a robust and re-writable epigenetic patterning mechanism. At a conceptual level, the LEAP model shows how epigenetics can be viewed as an analog memory system. This perspective follows from a long tradition of understanding chromatin regulation as a form of neural computation (*38, 39*). A recent trend has been towards theoretically modeling chromatin in terms of energy landscapes; (*40*) suggested that reprogramming can be thought of as annealing on a chromatin energy landscape, (*41*) connect liquid-liquid phase condensates to programmable spatial Hopfield networks, and (*42, 43*) interpret enzyme-mediated polymer looping as an energy-based generative model. The LEAP model stands out from these works because it moves beyond theoretical frameworks to model specific biological molecules and epigenetic marks. In the process, we find that cells can implement a flexible computational system capable of maintaining a wide range of patterns without specifying an external (or learned) energy function or landscape.

Additionally, the LEAP model makes several qualitative predictions which agree with previous literature. First, the LEAP model predicts that the total amount of methylation is approximately conserved (see SI F) which is consistent with experimentally observed histone mark redistribution (*44–47*). This behavior follows from the limited pool of methyltransferase enzymes in our model; the stabilizing role of enzyme limitation in setting the total level of modification was first developed in physical models of epigenetic maintenance (*17*) and subsequently generalized as enzyme titration (*15, 18*). This can be seen in the sensitivities of epigenetic patterns to the amount of enzyme *E* and the division time which conceptually control the effective size and density of epigenetic regions as depicted in Figure 2 G and H. The sensitivity analysis suggests that faster dividing cells must either have higher levels of enzymes to compensate, or lower levels of the methylation mark. Quantitative measurements of PRC2 proteins in different cell types show an approximately tenfold higher number of enzymes in mouse embryonic stem cells, which are reported to divide every ∼ 4 − 12 hours (*48, 49*) compared to both glioma and human embryonic kidney cells (HEK), which divide roughly every 48 hours (*50*). Comparing glioma cells with HEK cells, which have roughly equal doubling times, HEK cells have about two times more enzymes than glioma cells, and also a higher percentage of methylated histones (*50*). Our model is also consistent with the findings of Jadhav et al. (*51*), who experimentally decreased the levels of methyltransferase, leading to replicative dilution of the mark; in addition they observed that dilution per cell cycle is dependent on replication speed, with faster dividing cells diluting faster.

We demonstrate the generalizability of our model via genome-wide histone methylation mechanistic simulation using data from two cell types, two methylation marks, and 8 different donors. The correlations between the simulations and data indicate that the LEAP model can reliably explain epigenetic maintenance at the TAD scale. However, the LEAP model performs considerably worse on small length scales and tends to over-estimate epigenetic drift when compared to cell-passage experiments. Jointly these results indicate that there are additional biological processes biasing the formation of and maintenance of epigenetic patterns.

A central implication of these results is biological. Because LEAP reproduces genome-wide inheritance of H3K9me3 and H3K27me3 using only polymer looping and enzyme co-localization, a baseline of epigenetic memory does not require the specialized elements that many models invoke, such as dedicated nucleation sites, boundary or insulator elements, mutually antagonistic marks, or large-scale chromatin compaction. This does not imply that such elements are absent from the genome; nucleation sites (*52*) and boundary elements (*53*) are well documented at particular loci. Rather, our results indicate that these features are not universally required wherever epigenetic memory occurs, and need not be present in equal measure at every epigenetic domain. Consistent with this view, heterochromatin can be inherited through many cell divisions after removal of the sequence-specific nucleator (*54, 55*), modified histones alone can template inheritance (*56*), and Polycomb domains are frequently found in a decompact rather than collapsed state (*57*). We therefore interpret LEAP as a default, genome-general mechanism of epigenetic maintenance, upon which locus-specific elements and additional biological processes act to refine, stabilize, and regulate the resulting patterns.

Finally, we capitalize on this framework model to build a simple model to mimic epigenetic rejuvenation that captures the capability of metazoan cells to be reprogrammed. Our simulations demonstrate a mechanism by which epigenetic patterns can persist after the reprogramming signal has been removed. In this model, a bias signal promotes methylation in the regions overlapping with the original pattern. This is consistent with the idea that pioneering transcription factors, such as the Yamanaka factors, when dosed and timed appropriately can reactivate the developmental programs that established the somatic patterns (*58–60*). The model can also be instructive about chemical reprogramming based on the observation that small molecules may disrupt chromatin phase condensates (*61*). Rejuvenation in our model arises naturally through a dynamic reorganization of the epigenome caused by accumulation of epigenetic modifying enzymes in regions becoming increasingly methylated through the action of the bias signal. This causes regions to which epigenetic marks have drifted to slowly lose their marks through a competitive process similar to transcription factor theft (*62*), without a need to recognize, keep track of, or know *a priori* in any way the genomic location of epigenetic drift. A consequence of this fact is the observation that reprogramming can occur in many different cell types. This hints at the possibility of partial reprogramming exploiting the precise regulation afforded by the co-evolution of transcription factors, epigenetic regulators, and the physical substrate they bind to during development.

## Data Availability

Data are available on GEO with Accession Number GSE320278.

## Supplemental Information

### A Idealized LEAP Model

Let us consider a simplified epigenetic code consisting of a single epigenetic mark which resides along the monomers of a polymer chain representing chromatin. This polymer has *N* monomers (nucleosomes containing a modifiable histone), each denoted by *x_i_* ∈ {0, 1}, which indicates whether the histone *i* is unmodified (*x_i_* = 0) or modified (*x_i_* = 1). The ordering of these monomers in a linear chain *X* = (*x*_1_, …, *x_i_*, …, *x_N_*) constitutes a specific epigenetic pattern.

The process of epigenetic modification occurs through the action of enzymes capable of both reading and writing biochemical modifications on histones (*63*). A common property of reader/writer enzymes is that they “spread” modifications from modified to unmodified histones that are co-localized (*63*). For this to happen an unmodified nucleosome has to come into contact with a modified nucleosome, and a reader/writer enzyme has to be present to catalyze the reaction. We postulate that epigenetic pattern restoration is rooted in two physical properties: 1) an inverse power law decay with separation distance in chromatin contact probabilities; and, 2) the ability for epigenetic reader/writer enzymes to co-localize with their own reaction product. We now turn our attention to these two requirements.

#### Contact Probability

We first seek the probability that an unmodified nucleosome comes into contact with a modified nucleosome. Chromatin is at thermal equilibrium with the rest of the cell which gives rise to Brownian motion (*64*). This diffusion driven motion of the polymer chain allows the polymer to explore its conformational space, effectively sampling the probability distribution that one chromatin locus comes into contact with another chromatin locus. The entropic cost of bringing together two monomers on a polymer is a logarithmic function of the number of monomers in the loop. By combining the Gibbs free energy change associated with this change in entropy with the Boltzmann distribution, a scaling relationship between the probability of loop formation and the length of a loop is obtained. It is well established that polymer contact probabilities decay with separation distance as an inverse power law (*28*):

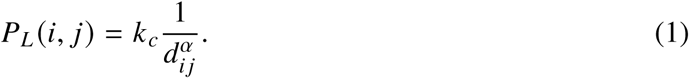

**Table S1:** Comparison of models of epigenetic memory. Model type, ingredients, and data validation for representative models of epigenetic memory, with the **LEAP** model (this work) in the final column. ✓ = used; (✓) = present implicitly; ∼ = partial, optional, or qualitative; – = not used. *Statistical contact law*: LEAP, Dodd, Erdel, Nickels, and Abdulla impose an ideal-polymer power law (∼ 1/*d^α^*), Sandholtz (Biophys. J.) uses a structure-dependent wormlike-chain looping law, and (✓) denotes contacts that emerge implicitly from an explicit coordinate polymer. *Explicit 3D conformation* denotes coordinate Monte-Carlo/molecular-dynamics polymer models. Hathaway’s model is steady-state (multi-generation memory shown experimentally, not simulated). *Validation data (modality; organism):* Hathaway, ChIP-seq (mouse); Erdel, ChIP-seq/qPCR reanalysis (fission yeast); Dodd, gene-silencing switching rate (fission yeast); Nickels, qualitative ChIP profiles (yeast); Abdulla, ChIP-seq (*γ*H2AX in human, H3K9me in mouse, reporter in fission yeast); Palacios, reporter assay (hamster, CHO-K1); Sandholtz-BJ, ChIP-seq (mouse); Sandholtz-PNAS, ChIP-seq and Hi-C (human); Owen, no dataset (qualitative correspondence); LEAP, CUT&RUN (human). For *genome-wide*, ∼ denotes a single chromosome (Sandholtz-PNAS) or genome-wide domain-shape statistics from a single-domain model (Hathaway). *Marks:* LEAP, H3K9me3 and H3K27me3; Palacios, DNA methylation, H3K9me3, and H3K4me3; Abdulla, *γ*H2AX and H3K9me2/3 in separate systems. *Donors/cell types:* LEAP, 8 HDF donors across HDF and HUVEC; Hathaway, ESC and MEF. Abbreviations: Hath’12 (Hathaway et al.), Erd’16 (Erdel & Greene), Dodd’07, Nick’23 (Nickels & Sneppen), Abd’22 (Abdulla et al.), Pal’25 (Palacios et al.), San-BJ (Sandholtz et al., Biophys. J.), San-PN (Sandholtz et al., PNAS), Owen’23 (Owen et al.).

|  | Hath’12 | Erd’16 | Dodd’07 | Nick’23 | Abd’22 | Pal’25 | San-BJ | San-PN | Owen’23 | **LEAP** |
|---|---|---|---|---|---|---|---|---|---|---|
| **Model type** |  |  |  |  |  |  |  |  |  |  |
| Formation / spreading | ✓ | ✓ | ✓ | ✓ | ✓ | ✓ | ✓ | ~ | ✓ | ✓ |
| Maintenance across divisions | ~ | ✓ | ✓ | ✓ | ✓ | ✓ | – | ✓ | ✓ | ✓ |
| Reprogramming / re-writability | – | – | – | – | – | ~ | – | – | – | ✓ |
| **Model ingredients** |  |  |  |  |  |  |  |  |  |  |
| Nucleation site | ✓ | ✓ | – | ✓ | ~ | – | ✓ | – | – | **–** |
| Boundary / insulator | – | ~ | ✓ | ✓ | – | – | ~ | – | – | **–** |
| Locus-specific parameters | – | – | – | – | ✓ | – | ~ | – | – | **–** |
| Antagonistic marks | – | ~ | ✓ | ✓ | ~ | ✓ | – | – | – | **–** |
| Histone–DNAme cross-talk | – | – | – | – | – | ✓ | – | – | – | **–** |
| Statistical contact law | – | ✓ | ✓ | ✓ | ✓ | – | ✓ | (✓) | (✓) | ✓ |
| Explicit 3D conformation | – | – | – | – | ~ | – | ~ | ✓ | ✓ | **–** |
| Enzyme co-localization / limitation | – | – | – | – | ✓ | – | – | ✓ | ✓ | ✓ |
| Compaction / self-attraction | – | – | – | – | ✓ | – | – | ✓ | ✓ | **–** |
| **Model validation** |  |  |  |  |  |  |  |  |  |  |
| Validated qualitatively with real epigenetic data | ✓ | ✓ | ~ | ~ | ✓ | ✓ | ✓ | ✓ | ~ | ✓ |
| Validated quantitatively with epigenetic data | ✓ | ~ | – | – | ✓ | ✓ | ✓ | ✓ | – | ✓ |
| Validated on genome-wide epigenetic data | ~ | – | – | – | – | – | – | ~ | – | ✓ |
| Validated with multiple epigenetic marks | – | – | – | – | ~ | ✓ | – | – | – | ✓ |
| Validated with multiple donors | – | – | – | – | – | – | – | – | – | ✓ |
| Validated with multiple cell types | ✓ | – | – | – | ~ | – | – | – | – | ✓ |

Here, *P_L_* (*i*, *j*) is the contact probability of two polymer loci, *i* and *j*, *d_i j_* = |*i* − *j* | is the linear separation distance between the loci, *k_c_* is the contact probability rate related to the diffusivity of the chain and *α* is a scaling exponent. Experiments measuring chromatin contact probabilities genome-wide have confirmed this behavior for a wide range of length scales (*24, 29*).

#### Enzyme Localization

Next, we seek the probability that an enzyme is present at locus *j* to catalyze the modification of locus *i*. Experimental observations (*21, 23*) have shown that enzyme binding is preferential for already modified histones. First, we compute the concentration of bound enzymes as a function of the total available enzymes *E* = *E _f_* + *E_b_* and total monomers *N* = *N_M_* + *N_U_* where *E _f_*, *E_b_*, *N_M_* and *N_U_* denote the free enzymes, bound enzymes, number of modified monomers and number of unmodified monomers, respectively. In the limit of large *N_M_* and large *E*, we model the binding of an enzyme to any modified monomer as an ordinary differential equation:

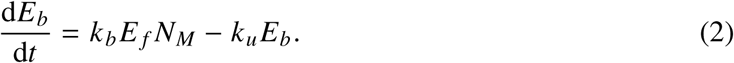

Here, *k_b_* is the binding rate of a free enzyme to a modified monomer and *k_u_* is the unbinding rate. We next solve for the steady state solution assuming that binding and unbinding reactions occur quickly:

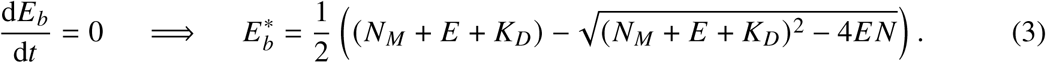

Here, 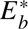 is the steady state concentration of bound enzymes and 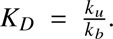 Additionally, we note that the binding curve between the catalyzing enzyme and modified histones has a sigmoidal shape suggestive of cooperative binding (*23*). We conjecture that this effect is due to preferential accumulation of enzymes at densely modified regions in the 3D polymer chain conformation, which we model phenomenologically with a Hill function of the local modified monomer density:

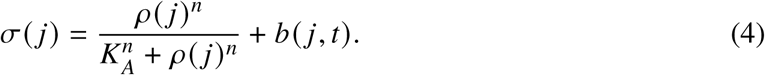

Here *ρ*(*j*) is the local density of modified monomer around *j*, *n* is a Hill coefficient representing the non-linear nature of enzyme localization, and *K_A_* is the histone methylation density that produces half max of the enzyme count. The term *b*(*j*, *t*) ≥ 0 is a position and time dependent bias term which can be used to model sequence dependence or introduce external perturbations; however, unless otherwise noted, we assume *b*(*j*, *t*) = 0. We chose to make this term additive (as opposed to multiplicative) because there is substantial evidence that epigenetic remodeling factors can act on both methylated and unmethylated regions (*65*). For example, Yamanaka factor signaling networks are known to target euchromatin and heterochromatin (*66*), while H3K9 and H3K27 methylation occurs only in heterochromatin. Therefore, we allow a bias which is independent of the methylation density. Combining equations (3) and (4) we arrive at an expression for the probability of an enzyme at monomer *j* :

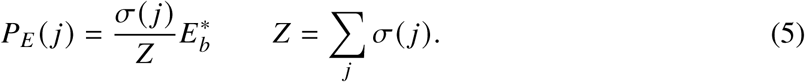

Note that normalizing by *Z* is required to ensure the correct total number of bound enzymes: 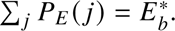

#### Continuous Time Markov Chain

Next, we model the process of epigenetic mark restoration. In this process, marks are stochastically added one at a time to unmarked monomers 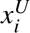 after they come into contact with an enzyme bound to marked monomer 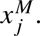 These microscopic spreading rates are given by:

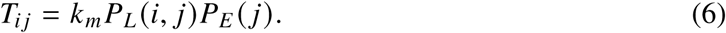

Here, *k_m_* is the enzymatic modification rate. Finally, we combine all the microscopic spreading rates into a continuous time Markov chain (CTMC) (*67*) representing a stochastic process of epigenetic mark restoration between cell divisions. The transition rates from state *X* to *X*^′^ are given by:

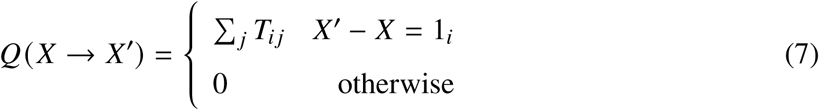

Here, *T_i j_* are the spreading events where an enzyme at *j* catalyzes the addition of a mark at *i*, and *X*^′^ − *X* = 1*_i_* denotes that the difference between *X*^′^ and *X* is the addition of a single modification at monomer *i*. This transition rate matrix governs the dynamics of a probability distribution of the epigenetic pattern P(*X*): <u>dP(</u> *<u>X</u>*) d*t*

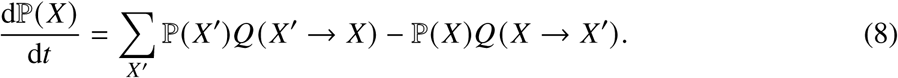

#### Computational Implementation

We begin each simulation as a sequence *X* that represents a particular epigenetic pattern. This sequence is used as the initial state in the CTMC represented by equation (7). This CTMC is simulated for a fixed amount of time, akin to cell division time, *t_div_* via the *τ*-leaping method (*68*). The density *ρ*(*j*) in equation (4) is approximated as a Gaussian kernel density estimate along the polymer chain, evaluated at monomer *j* :

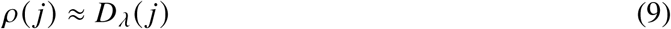

where *D_λ_* is a one-dimensional Gaussian kernel density estimate with bandwidth *λ*. This estimate is updated every iteration of the *τ*-leaping algorithm.

When *t_div_* is reached, the cell divides resulting in the stochastic partitioning of all epigenetic marks randomly between the two daughter cells. Because only a single cell is tracked in our simulation, this amounts to randomly erasing each epigenetic mark with a probability of 50%. The resulting post division state, *X*^′^, is then used as the initial state that can be simulated as a CTMC until the next division.

#### Model Parameterization

We used the following parameters for our simulations which are estimated from literature. The number of monomers, *N*, is set to 10000 monomers as a representative sequence length. The number of histone methyltransferases, *E*, varies widely by cell type and organism, but is always one or more orders of magnitude less than the number of histones. Therefore, we choose *E* = 1000 enzymes (*50*). The methyltransferase-histone dissociation constant was estimated to be *K_D_* = 300 (which is in units of molecular counts) based on (*21*). Measurements for the enzymatic rate constant vary by orders of magnitude, from 10^−3^ as measured in (*69*) to 30 as measured in (*70*) and we used the intermediate value of *k*_meth_ = 1.0 (in units of per day). The division time of a eukaryotic cell ranges from about a day for rapidly dividing cells to years for slowly dividing cells, but experiments are typically conducted on rapidly dividing cells to shorten the length of experiments. We therefore choose a division time of 1 day. The density bandwidth *λ* measures the length scale over which nearby nucleosomes compact and should be roughly the persistence length of the nucleosome chain, which is approximately 10 nucleosomes (*71*). Finally, we estimated the Hill function parameters *K_A_* = 0.00025 (which is in units of monomer density along the sequence) and *n* = 3.3 from the binding curve for the H3K9me3 methyltransferase SUV39H1 measured in (*21*).

### B Spontaneous Pattern Formation

In this section, we investigate some properties of the idealized LEAP model and discuss how they give rise to spontaneous pattern formation.

We begin by briefly commenting on some of the mathematical properties of this model. First, this model is trivially ergodic - meaning any state can in theory be reached from any other state - due to the ability to add individual marks at any location and then partition the marks randomly into two arbitrary new states (provided no new marks are created). Second, the dynamics of this system are monomer location invariant (neglecting edge effects of the chain) provided that the bias term is 0. Jointly, these properties suggest that the steady state of a very large *N* idealized model should be almost uniform.

These observations are not at odds with the fact that the LEAP model maintains patterns. Indeed, steady state dynamics of a CTMC are defined as an infinite time average which does not imply that the short term dynamics of any given trajectory should appear uniform. We can observe aspects of this behavior via numerical experiments. In Figure 5 we show a sampling of idealized model simulations from random initial conditions. Fluctuations in initial condition density coupled with the stochastic nature of the model reliably give rise to persistent banded structures, a form of spontaneous pattern generation. Indeed, a fundamental hypothesis implicit in our model is that cells make use of the spontaneous pattern formation and restoration properties intrinsic in looping, enzymes, and phase condensates as a flexible and modulatable epigenetic memory system. While beyond the scope of this paper, we note that the ability to form epigenetic patterns spontaneously has potential implications for epigenetic evolution.

**Figure 5:**
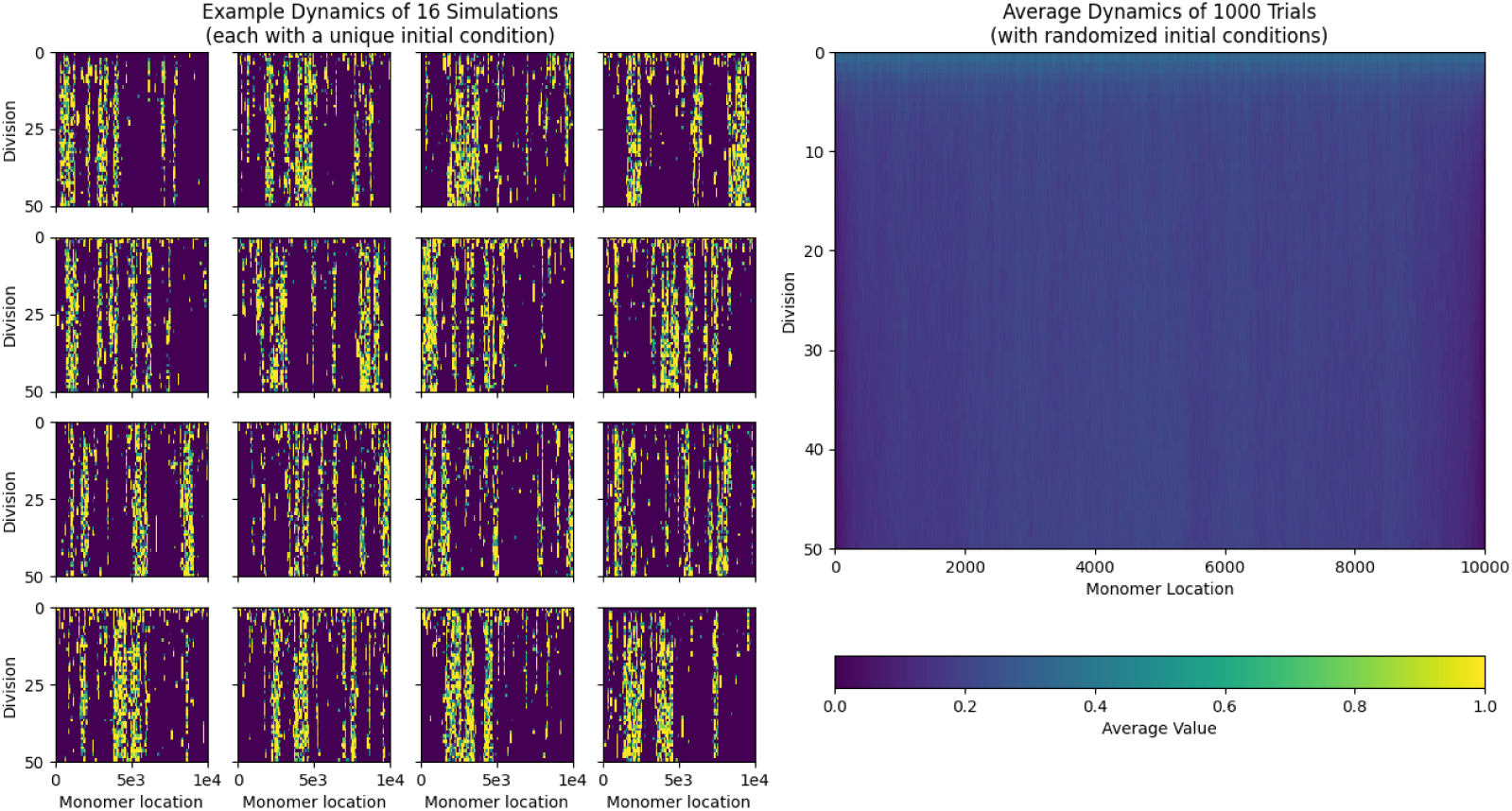
Left. Example simulations with randomized initial conditions spontaneously form banded patterns. Right. Average of 1000 randomized simulations indicates that the steady state distribution is uniform.

### C Coarse Grained LEAP Model

In this section, we describe the coarse grained model used for genome-scale epigenetic simulations. In addition to the coarse-graining approach, this model also involves multiple chromosomes and several assumptions had to be made. First, we (very approximately) assume that the chromosomes do not interact in 3D space, meaning there is no contact probability between different chromosomes. Although this is obviously not accurate, it is a necessary approximation for our simple model to be applicable. However, we do couple the chromosomes via a shared pool of enzymes. Specifically, we partition the total enzymes *E* between the different chromosomes in a way proportional to the 3D volume occupied by each chromosome. In the following section, we will derive the coarse grained model from the un-coarse grained *detailed* model. Note that due to having multiple chromosomes, the notation used in this section is slightly different from the rest of the paper.

The detailed state *X_c_* of a chromosome *c* with *N_c_* monomers each with 0 or 1 marks is denoted by:

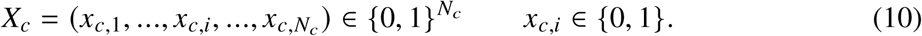

We coarse grain the detailed polymer *X_c_* into a shorter polymer *Y_c_* of length *ν_c_* = *N_c_*/*μ* where each monomer may have up to *μ* marks. In other words, the coarse graining procedure combines *μ* binary monomers in *X_c_* into a single coarse-grained monomer in *Y* which may have up to *μ* marks. Denote the coarse grained polymer:

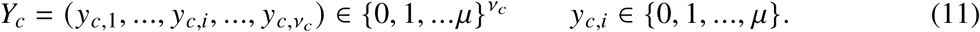

The value of each individual coarse grained monomer *y_c_*_,*i*_ is a sum of monomers from the detailed state *x_c_*_,*i*_:

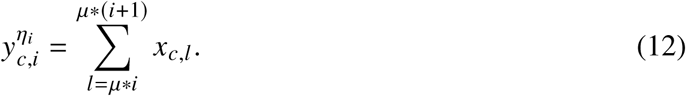

Next, coarse grained chromosomes are coupled via a shared pool of enzymes partitioned proportional to their occupied volume *V_c_*:

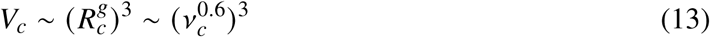

Where 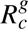 is the radius of gyration of chromosome *c* and the final scaling law relates the radius of gyration to the number of monomers in a self-avoiding polymer chain (*72*). Using the result from equation (3), the total number of bound enzymes to chromosome *c*, *E_c_*, is given by:

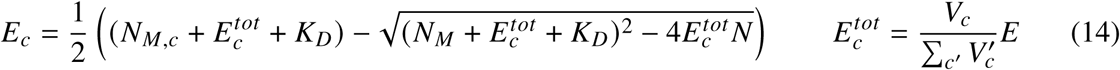

Here, *N_M_*_,*c*_ is the number of marked monomers in a chromosome *c* and 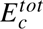 is the total number of enzymes available to chromosome *c*.

We now have the necessary pieces to write exact transition rates between coarse grained states *Y* and *Y*^′^:

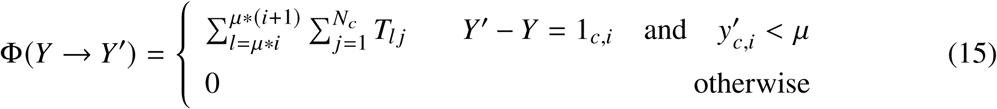

Here, transitions add a single mark to a coarse-grained monomer at a time which has at most *μ* marks. The term 1*_c_*_,*i*_ indicates that *Y* and *Y*^′^ differ by the addition of a single mark at chromosome *c* monomer *i*. The double sum first sums over the monomers that are combined and then over the transition rates of those individual monomers as defined in (6). Next, in order to efficiently simulate at the coarse-grained scale, the double sum must be approximated in terms of the coarse-grained polymer:

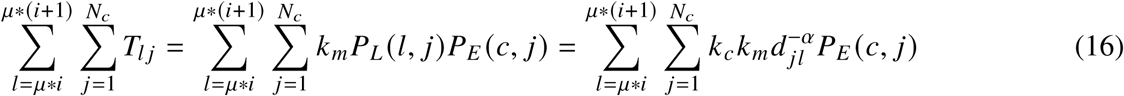

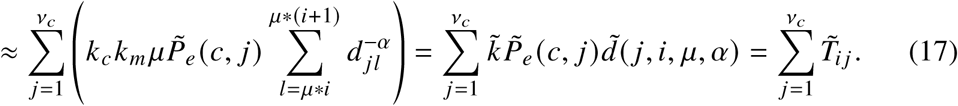

Where

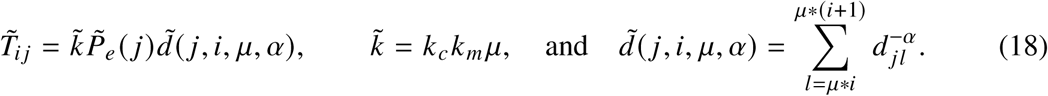

Here, we have rewritten the sum over all monomers from the detailed state into a sum over coarse grained monomers. Two approximations were made to get this result. First, the probability of enzyme localization *P_E_* (*c*, *j*) of the detailed model is approximated by the probability of enzyme localization of the coarse-grained monomer, *P̃_e_*(*c*, *j*). Inspired by the renormalization group theory of the Ising model (*73*), this function is assumed to be constant across the set of monomers in the sum and have the same functional form as *P_E_* :

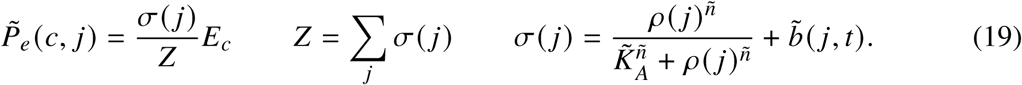

Here, *ρ*(*j*) is the marked monomer density at location *j*. Additionally, the sigmoid parameters *K̃_A_*, and *ñ* must be derived from data and are not necessarily the same as the detailed model’s parameters. Finally, we have let *b̃* (*j*, *t*) = 0 for simplicity. Second, the inverse power law is replaced by a surrogate function which adds up the contributions of the power law over each coarse-grained region of the polymer. Importantly, due to being density-independent, *d̃* can be pre-computed as a look-up table for computational efficiency. This allows us to write an approximate coarse-grained transition rate function Φ̃ (*Y* → *Y*^′^):

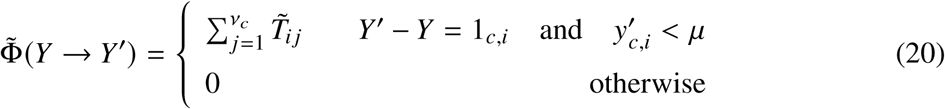

The CTMC for the coarse grained model will then follow the following master equation:

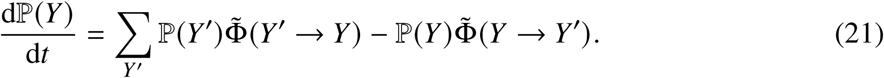

### D Full Genome Epigenetic Simulations

#### D.1 Full Genome Simulation Results

HDF full genome simulations were carried out by coarse-graining the HDF data (H3K9me3 and H3K27me3 CUT&RUN from 8 donors) and then simulating the entire genome of each donor for 200 generations after initializing the model to the coarse-grained data from that donor. Ten replicates were carried out per donor, resulting in a total of 160 full genome simulations. Large scale patterns (on the order of megabases) were faithfully maintained over the entire duration of the simulation. The average output from the 10 replicates of each full genome simulation and the coarse grained data is shown in Figures 6 and 7.

**Figure 6:**
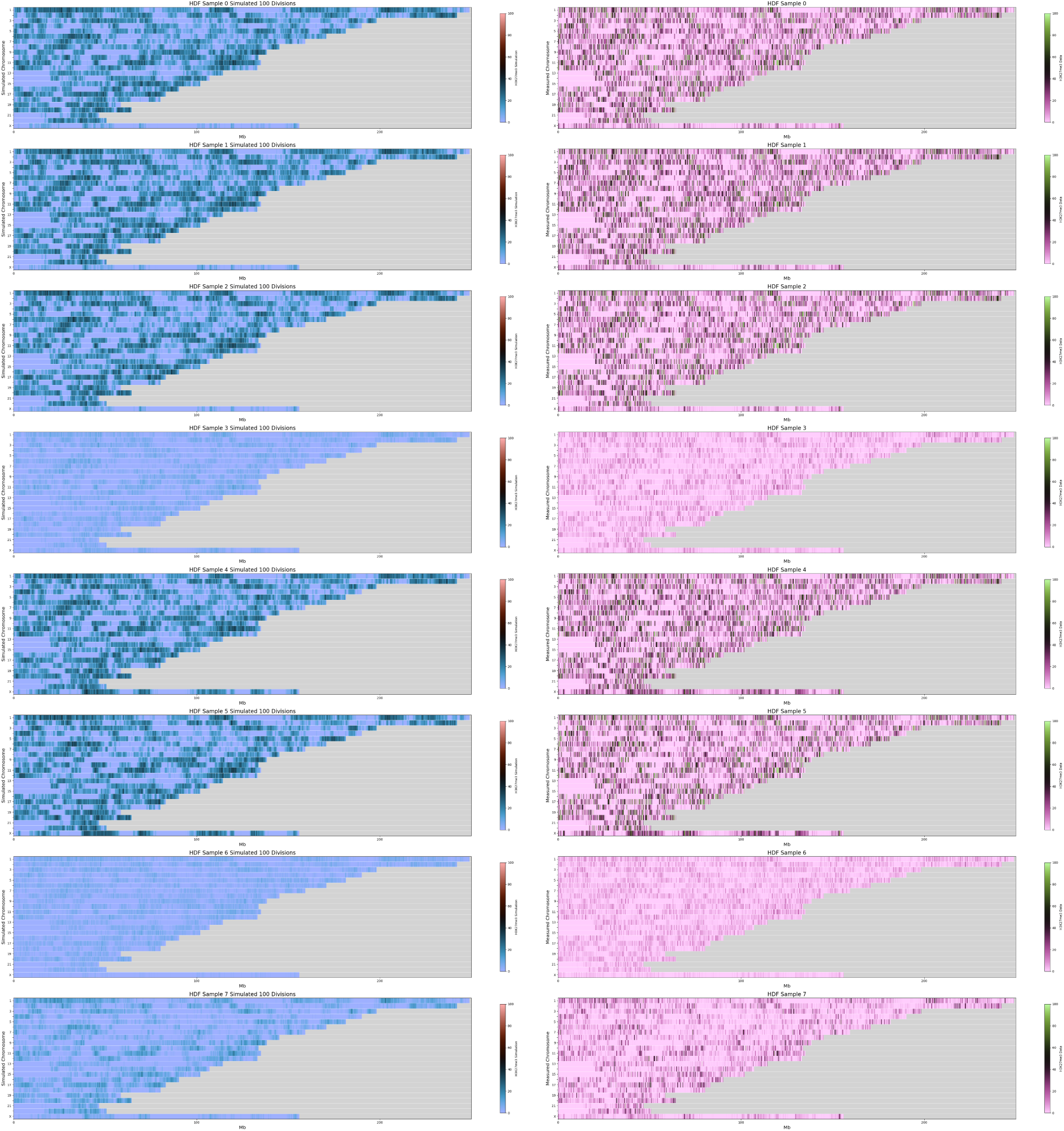
HDF H3K27me3 coarse simulation (blue heatmaps) results after 100 divisions and coarse-grained data (pink heatmaps) by donor and chromosome. Not smoothed.

**Figure 7:**
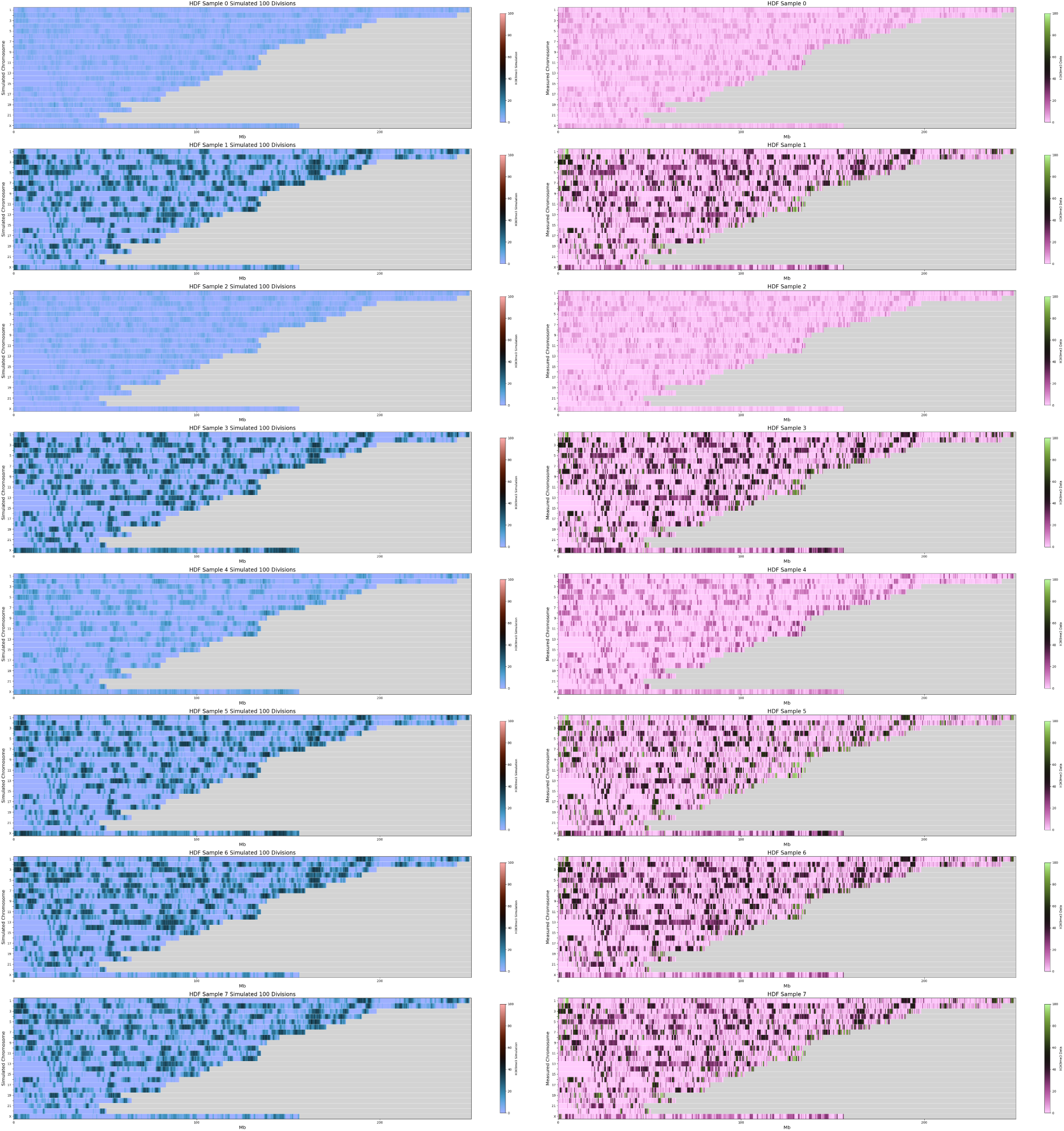
HDF H3K9me3 coarse simulation results (blue heatmaps) after 100 divisions and coarse-grained data (pink heatmaps) by donor and chromosome. Not smoothed.

**Figure 8:**
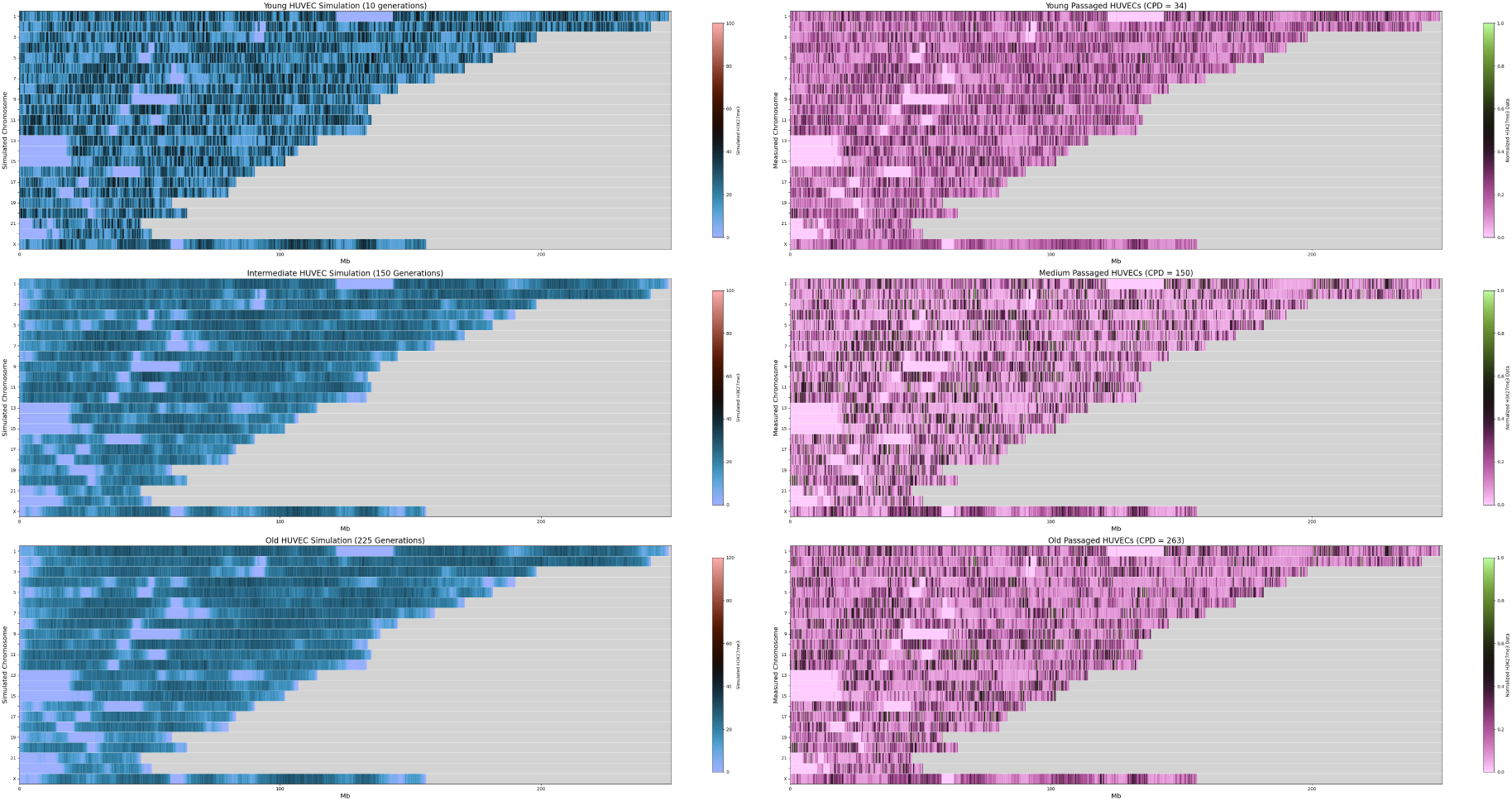
HUVEC H3K27me3 coarse grained data and simulation results at different passages and divisions. Not smoothed.

HUVEC full genome simulations were carried out analogously to the HDF simulations by initializing the LEAP model to the young passaged HUVEC H3K27me3 data and simulating 25 replicates for 230 generations. The simulated epigenetic dynamics were then compared to coarse grained data from the medium and old passaged HUVECs. Although we see greater epigenetic drift in the model than in the data, the model is still able to accurately predict maintenance of large-scale epigenetic domains.

#### D.2 Coarse Graining Data

In order to compare the coarse grained model to real epigenetic data, it is necessary to coarse-grain the bulk epigenetic data in an analogous way to the model. This procedure is also used to generate initial conditions for the coarse-grained model from data. Briefly, raw reads from the CUT&RUN experiment are aligned to a reference genome and converted to counts per nucleotide as described in SI G.4. These data are effectively noisy and unitless quantities proportional to the number of marked histones at any given genomic location across a large population of cells. First, we bin and compress the raw underlying data *Z* to compute a one-normalized local density *ρ̄*:

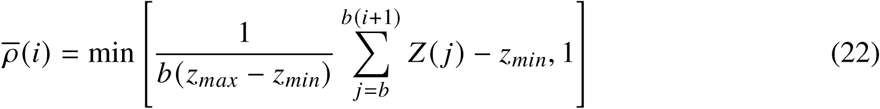

Here, *b* is the bin-size, chosen to match the resolution of the coarse grained model, and *z_max_* and *z_min_* are empirically determined parameters. We found that *z_max_* = 4 and *z_min_* = 0 gave reasonable results for our data set. In particular, *z_max_* = 4 was chosen to compress the extreme values found in blacklist regions (*74*) as we cannot discard these regions of each chromosome when running mechanistic simulations.

#### D.3 Generating Initial Conditions from Data

We generate initial conditions 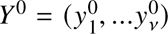 by sampling a binomial distribution 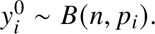 Here, *n* = *μ* is the maximum number of marks per coarse-grained monomer and *p_i_* = *ρ̄*(*i*) is the probability of any particular site in the coarse-grained monomer being occupied.

#### D.4 Coarse CTMC Simulation Details

We simulate the coarse grained CTMC using the *τ*-leaping algorithm (*75*) with *E_c_* and *T̃* recomputed at each *τ*-step. Analogously to the idealized model, marks are partitioned at regular cell divisions based upon the binomial distribution.

All simulations were run using custom Python code on 48-CPU AWS nodes running Linux with 384 GB of RAM. Simulations were run sequentially on any given node but were accelerated due to the automatic parallelization of NumPy array operations.

We were able to run hundreds of full genome simulations using just 4 nodes in a little over a week. For each of 8 donors, 10 replicates of H3K9me3 and H3K27me3 epigenetic models were simulated for 200 cell divisions starting at initial conditions derived from CUT&RUN data. Additionally, 20 replicates of HUVEC H3K27me3 simulations were conducted for 230 divisions with initial conditions derived from the young (low passage) HUVEC samples.

#### D.5 Coarse CTMC Simulation Parameterization

The parameters used for the coarse-grained simulations are as follows. The bin size was set to 10, 000 base-pairs, which amounts to coarse graining the model by a factor of ∼50 nucleosomes (200 bp per nucleosome). *z_min_* and *z_max_* were set to 0.0 and 4.0 respectively, which we found reduced the incidence of problematic peaks in the signal, often corresponding to blacklisted CUT&RUN regions (*74*). The sigmoid parameter *K_A_* was set to 3.5 × 10^−7^ (in units of methylation density per coarse-grained monomer), *n* in the sigmoid was set to 3.3 as in the idealized model. *λ*, the bandwidth used in the density estimate, was set to 10, which in the coarse grained model amounts to 100 kb in genomic space. The methyltransferase-histone dissociation constant *K_D_* was set to 200 molecules. The division time *t_d_* was set to 1.667 days which is the average time between divisions for our HUVEC cells. Our timestep was set to 0.667 days, which amounts to 10 timesteps per division and was chosen to both maintain accuracy and minimize computational requirements. *α*, the inverse power law decay exponent, was set to 1.5 as in the idealized model and falls within experimentally measured values of *α* (*24, 29*). The total number of enzymes across all chromosomes was set to *E* = *N* × bin size/(base pairs per mark × 100) which amounts to choosing *E* to be 100 times less than the number of histone substrates. Lastly, we use a methylation rate *k*_meth_ of 0.001657 per day for the HDF cell simulations, and a value of 0.001653 per day for the HUVEC cell simulations.

**Figure 9:**
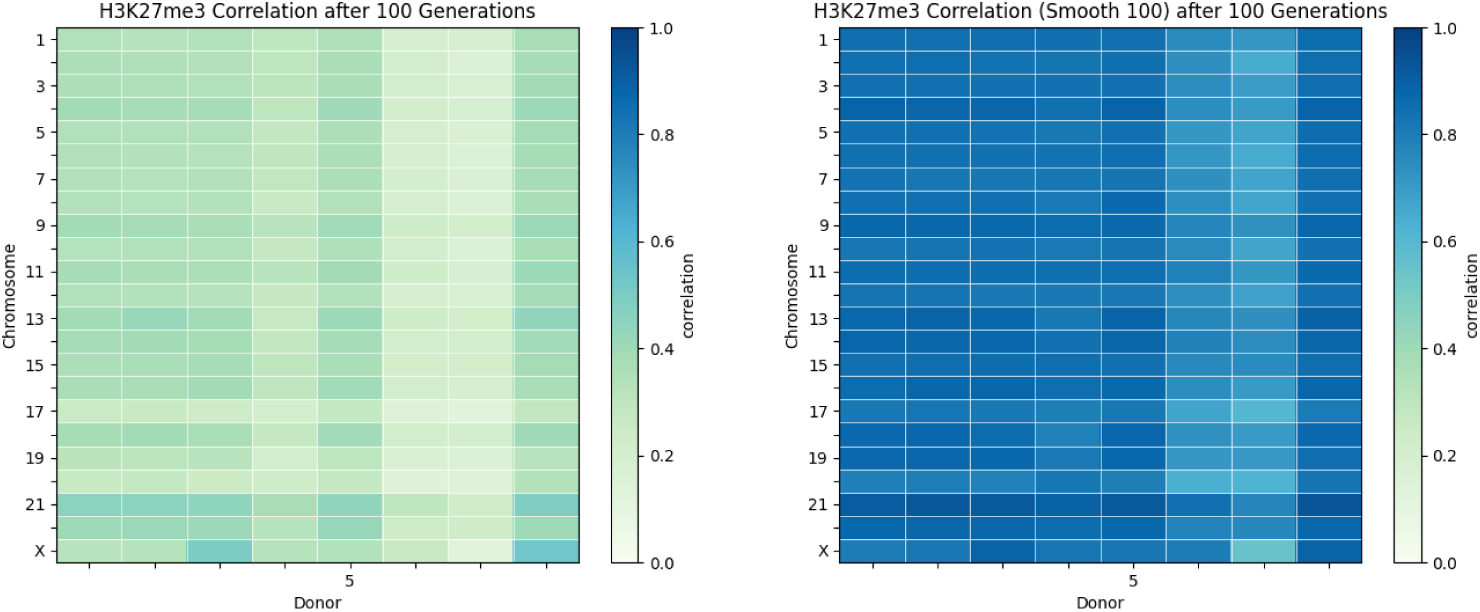
Correlations between coarse-grained data and simulation results (after 100 divisions) by donor and chromosome for smoothed and un-smoothed H3K27me3.

Importantly, we note that by measuring histone marks along the genome using CUT&RUN, we do not necessarily know the true amount of the mark at each genomic location, but rather the number of reads that were mapped to each genomic location. It is thus likely that those parameters in our model that interact to give rise to a total methylation are somewhat disconnected from the true biological values. We hypothesize that this is what allowed us to use almost identical parameter sets in both cell types, both types of marks, and for all donors.

### E Correlations and Autocorrelation Analyses

#### E.1 Computing Correlations

Correlations between a simulation state and data were computed using SciPy’s implementation of Pearson’s correlation between a simulated epigenetic state and the one-normalized density *ρ*(*i*).

Correlations between smoothed states were computed using a Pearson correlation on smoothed epigenetic states and smoothed one-normalized densities. Smoothing was carried out with a 1-dimensional uniform smoothing using window sizes of 100 unless otherwise noted.

Note that we excluded chromosome Y from these analyses because in the majority of simulations it lost all methylation and we therefore could not compute correlations.

**Figure 10:**
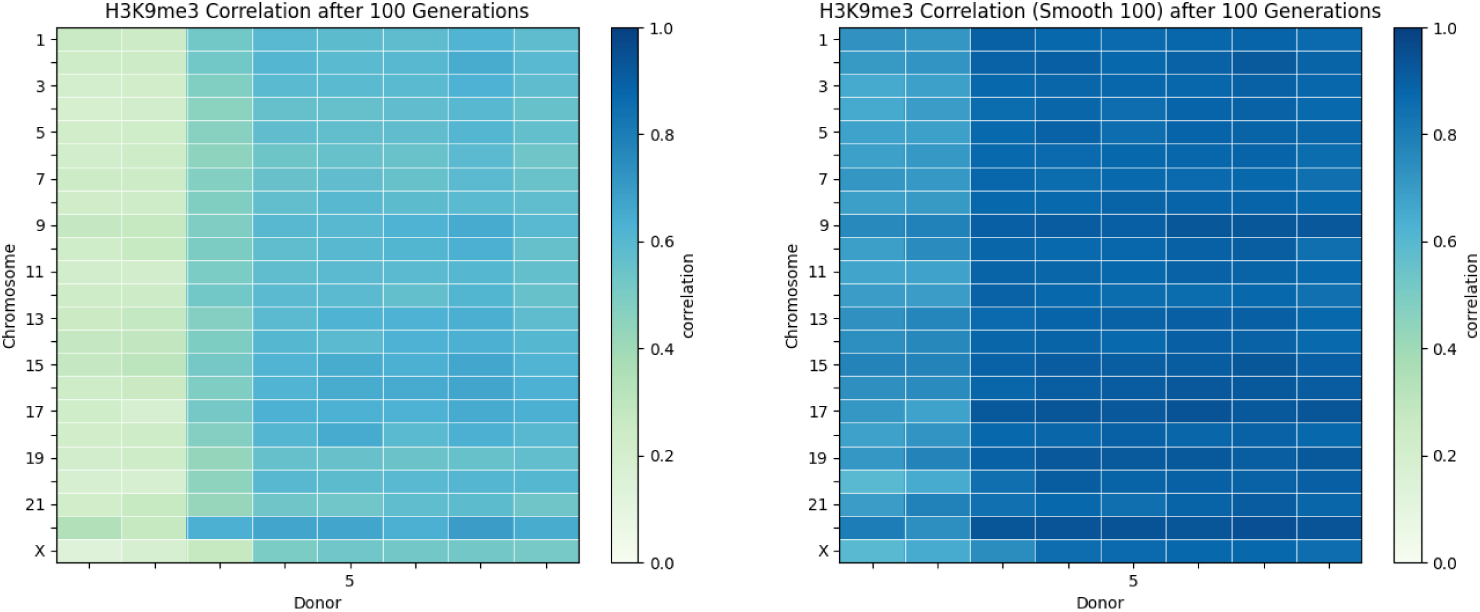
Correlations between coarse-grained data and simulation results (after 100 divisions) by donor and chromosome for smoothed and un-smoothed H3K9me3.

#### E.2 Computing Auto-correlations

Auto-correlation functions of epigenetic simulations were computed numerically by computing the Pearson correlation between the initial state and subsequent states at each time point *t* averaged together across multiple replicates (in the case of HUVECs) and replicates and donors (in the case of HDFs).

Auto-correlation functions of smoothed epigenetic simulations were computed numerically by computing the Pearson correlation between the smoothed initial state and subsequent smoothed states at each time point *t* averaged together across multiple replicates (in the case of HUVECs) and replicates and donors (in the case of HDFs). Smoothing was carried out with a 1-dimensional uniform smoothing using window sizes of 100 unless otherwise noted.

### F Conservation of Epigenetic Marks Analysis

First we comment that the CTMC described in equation (8) does not explicitly include mark degradation; this was included as regular binomial partitioning in our simulations to better mimic cell division. However, our implementation of binomial partitioning would be complex to consider in full CTMC formalism. Thus, we first simplify the model and allow marks to degrade at a rate 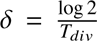 where *T_div_* is the time between cell divisions. This value of *δ* ensures that in *T_div_* of simulation, each mark has a fifty percent chance of being degraded, consistent with our previous model. Second, instead of considering the full distribution of epigenetic marks, we consider a simplified bulk model. In this model, the total number of marks is partitioned into marked and unmarked continuous variables which are governed by the following ODE:

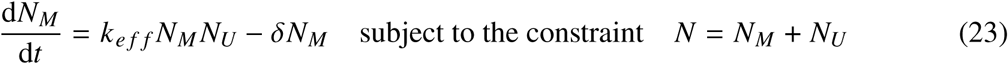

**Figure 11:**
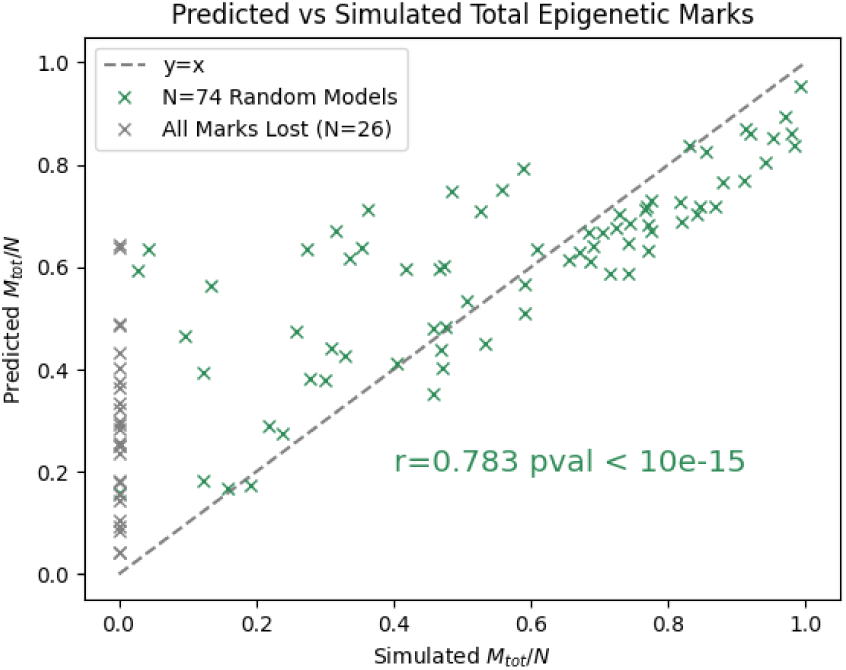
Steady state total epigenetic marks for randomly parameterized models compared to analytic prediction.

where *N* is the polymer length, *N_M_* is the number of marked monomers, *N_U_* is the number of unmarked monomers, and *k_e f f_* is an effective methylation rate. Solving for steady state results in:

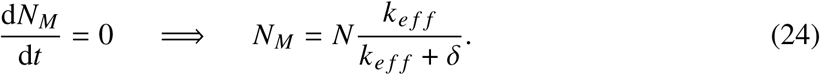

The effective methylation rate will be state dependent. Indeed, for a specific state *X*, it can be written exactly by summing over all possible methylation events:

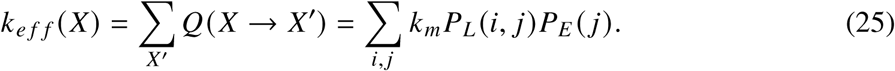

Both *P_E_* and *P_L_* depend on many details of the state *X* so we must approximate them. The contact probability *P_L_* (*i*, *j*) depends on the specific distance between the monomers *i* and *j*. We approximate this by assuming *i* is in the middle of the polymer and averaging over possible values of *j* :

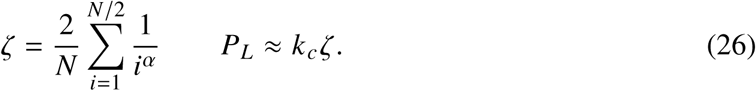

The bound enzyme probability depends on the total number of marked monomers. However, the total number of bound enzymes can never exceed *E*. Therefore, we bound Σ*_j_P_E_* (*j*) < *E*. Combining these approximations we arrive at:

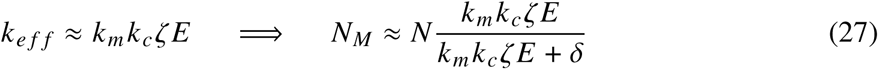

### G Experimental Methods

#### G.1 Cell preparation and culture

Human Dermal Fibroblast lines were purchased from the following vendors: 9F (Coriell, Catalogue No. GM00038), 12M (Coriell, Catalogue No. AG16409), 21F (Coriell, Catalogue No. AG09309), 22Ma (Coriell, Catalogue No. GM05294), 22Mb (Coriell, Catalogue No. GM23815), 28Fa (Cell Applications, Catalogue No. 106-05a, Lot No. 2146), 28Fb (Cell Applications, Catalogue No. 106-05a, Lot No. 2049), 29M (Coriell, Catalogue No. AG04054). Cells were maintained in a 37^◦^C hypoxic incubator (5% CO_2_, 5% O_2_) using DMEM/F-12 GlutaMAX media (Gibco, Catalogue No. 31331093) supplemented with 10% heat-inactivated Fetal Bovine Serum (Thermo Fisher, Catalogue No. 17593595), 1X MEM-NEAA (Gibco, Catalogue No. 11140035), 50 µM 2-Mercaptoethanol (Gibco, Catalogue No. 31350010), and 16 ng/mL FGF-2 (Gibco, Catalogue No. PHG0360).

For nuclei isolation, cells were seeded and harvested two days later. Briefly, medium was removed, cells were washed once with 1X DPBS (Gibco, Catalogue No. 14190169), and trypsinized using TrypLE Express (Gibco, Catalogue No. 12605010). After cells were detached, medium was added to quench the TrypLE Express solution. Cells were transferred to a 15 mL Eppendorf tube and washed once with 1X DPBS. Nuclei were extracted using freshly prepared Nuclei Extraction Buffer (20 mM HEPES pH 7.9, 10 mM KCl, 0.1% Triton 10X, 20% Glycerol, 1 mM MnCl_2_, 0.5 mM Spermidine, 1X Protease Inhibitor Cocktail) and CUTANA ChIC/CUT&RUN manufacturer recommendations (Epicypher; Catalogue No. 14-1048). 600,000 cells per CUT&RUN experiment were slow-frozen in an Eppendorf DNA LoBind tube (Eppendorf, Catalogue No. 0030108051) and stored in a -80^◦^C freezer.

Human umbilical vein endothelial cells (HUVECs; Cell Applications) were cultured in Endothelial Cell Growth Medium (Cell Applications, 211–500) at 37 ^◦^C in a humidified incubator with 5% CO_2_. Culture vessels were pre-coated with 0.001% fibronectin (Sigma-Aldrich, F0895) in HBSS (Gibco, 14175-053) for at least 1 h prior to seeding. The medium was replaced every other day. Cells were subcultured at 70–90% confluency using Trypsin-EDTA (Gibco, 15400054), neutralized with soybean trypsin inhibitor (Invitrogen, 17075-029), and reseeded at 5 × 10^5^ cells per T75 flask. Continued passaging induces epigenetic aging, consistent with prior reports linking DNA methylation changes to cellular aging (*76*). For CUT&RUN assays, HUVECs were harvested two days post-seeding, pelleted by centrifugation at 1200 rpm for 5 min at room temperature, and resuspended in cold freezing medium consisting of 10% DMSO (Sigma-Aldrich, D2650) and 90% FBS (Gibco, 10500064). Samples were cryopreserved using a controlled-rate freezing container at −80 ^◦^C for 24 h before transfer to liquid nitrogen storage. 250,000 cryopreserved HUVEC cells were used for each CUT&RUN reaction.

#### G.2 CUT&RUN

CUT&RUN was performed using the CUTANA ChIC/CUT&RUN Kit (Epicypher; Catalogue No. 14-1048) and manufacturer recommendations. Briefly, Wash Buffer, Cell Permeabilization Buffer, and Antibody Buffer, were made fresh on the day of the experiment and kept at 4^◦^C. Concanavalin A (ConA) Beads were washed twice with cold Bead Activation Buffer. After the last wash, activated ConA beads were resuspended in cold Bead Activation Buffer, distributed to 8-strip tubes, and placed on ice.

Cryopreserved HUVECs were thawed and washed twice with HBSS, followed by centrifugation at 600 × g for 3 min at room temperature. The cell pellet was then washed with room-temperature Wash Buffer and resuspended in Wash Buffer, before adding to ConA beads.

Human Dermal Fibroblast nuclei in Nuclei Extraction Buffer were thawed on ice and spun down. Supernatant was removed, nuclei were resuspended in freshly prepared Nuclei Extraction Buffer, and added to 8-strip tubes containing ConA bead mixture. Samples were incubated for 10 minutes at room temperature. Tubes were placed on a magnetic rack to allow slurry to clear, and supernatant was discarded. Nuclei bound to ConA beads were resuspended in cold Antibody Buffer and 0.5 µg of either H3K9me3 (Active Motif, Catalogue No. 39161) or H3K27me3 (EpiCypher,

Catalogue No. 13-0055) antibody was added. Samples were gently vortexed and spun down before being placed on a nutator at 4^◦^C overnight.

The next day, samples were washed twice with cold Cell Permeabilization Buffer on a magnetic rack. Tubes were taken off the magnetic rack and samples were resuspended in cold Cell Permeabilization Buffer. Subsequently, pAG-MNase was added to the mix and tubes were gently vortexed. Samples were then incubated at room temperature for 10 minutes and washed another two times with cold Cell Permeabilization Buffer. Samples were resuspended in cold Cell Permeabilization Buffer and calcium chloride was added to a final concentration of 2 mM. Tubes were incubated on a nutator for 2 hours at 4^◦^C. Afterwards, Stop Buffer was added and samples were incubated at 37^◦^C for 10 minutes. Tubes were placed on a magnetic rack, and supernatants of CUT&RUN enriched DNA were transferred to new 8-strip tubes. SPRIselect beads were added at a concentration of 1.4X and incubated at room temperature for 5 minutes. Tubes were placed on a magnetic rack and supernatant was removed. Two 85% EtOH washes were performed and SPRIselect beads were air-dried for 2 minutes at room temperature. 0.1X TE Buffer was used to elute the DNA and CUT&RUN purified DNA transferred to new 8-strip tubes. Samples were quantified using the High Sensitivity Qubit dsDNA Quantification Kit (Thermo Fisher, Catalogue No. Q32851) before being stored in a -20^◦^C freezer.

#### G.3 Library Preparation

CUT&RUN library preparation was performed using the CUTANA CUT&RUN Library Prep Kit (Epicypher; Catalogue No. 14-1001 and 14-1002) and manufacturer recommendations. In brief, samples and kit reagents were thawed on ice. 10 ng of CUT&RUN enriched DNA was transferred to new 8-strip tubes and volume was adjusted to 25 µL with 0.1X TE Buffer. Samples were End Repaired, and Adaptors for Illumina were ligated followed by U-Excision digestion. SPRIselect beads were added at a 1X concentration and incubated at room temperature for 5 minutes. Tubes were placed on a magnetic rack and supernatant was removed. Two 85% EtOH washes were performed and SPRIselect beads were air-dried for 2 minutes at room temperature. 0.1X TE Buffer was used to elute the DNA. Indexing PCR was performed by adding unique i7 and i5 Primer combinations to samples followed by Hot Start 2X PCR Master Mix. 14 PCR cycles were used for Human Dermal Fibroblast and 11 PCR cycles were used for HUVEC samples to amplify the End Repaired and Adaptor Ligated DNA. A final PCR cleanup was performed with SPRIselect beads added at a 1X concentration, followed by two 85% Ethanol washes and resuspension in 0.1X TE Buffer. Samples were quantified using the High Sensitivity Qubit dsDNA Quantification Kit (Thermo Fisher, Catalogue No. Q32851) and amplified DNA was visualized using a High Sensitivity D5000 ScreenTape Assay for TapeStation Systems (Agilent, Catalogue No. 5067-5592).

#### G.4 CUT&RUN Sequencing and Data Processing

CUT&RUN libraries for HUVECs were sequenced on an Illumina NovaSeq 6000 system on S1 flow cells in 2 × 50 bp mode. Libraries for HDFs were sequenced on an Illumina NextSeq 2000 system on P2 flow cells in 2 × 150 bp mode (H3K9me3) or on P3 flow cells in 2 × 50 bp mode (H3K27me3). CUT&RUN data were then processed using the nf-core/cutandrun pipeline (v3.2.2) (*77*) implemented in Nextflow (v24.04.4) (*78*). The pipeline was executed on paired-end sequencing data from H3K27me3 and H3K9me3 experiments performed on human dermal fibroblasts (HDFs) from donors of different ages, as well as H3K27me3 experiments on in vitro passaged HUVECs. Raw sequencing reads were initially assessed for quality using FastQC (v0.12.1) (*79*). Adapter trimming and quality control were performed using Trim Galore (v0.6.6) (*80*) with Cutadapt (v2.10) (*81*) for adapter removal. The trimmed reads were aligned to the human reference genome assembly GRCh38.p13 (GCA 000001405.28) using Bowtie2 (v2.4.4) (*82*) with default parameters. Post-alignment processing was performed using SAMtools (v1.17) (*83*) for sorting, indexing, and generating alignment statistics. Duplicate reads were marked and removed using Picard MarkDuplicates (v3.1.0) (*84*). Quality control metrics of the aligned data were assessed using multiple approaches: Preseq (v3.1.1) (*85*) was used for library complexity estimation, and deepTools (v3.5.1) (*86*) was employed for generating quality metrics including correlation plots, PCA, and fingerprint plots. Coverage tracks were generated using deepTools bamCoverage and converted to bigWig format using UCSC tools (v377/445) (*87*) for visualization.

## Notes

### Competing Interest Statement

Most authors on this manuscript work at Altos labs, a company devoted to developing reprogramming based therapies.

### Summary of Updates

This is a new version of the manuscript based upon reviews. Major updates are in the introduction and discussion around how our work relates to prior work in the field.

